# Distinct Clinical Phenotypes in KIF1A-Associated Neurological Disorders Result from Different Amino Acid Substitutions at the Same Residue in KIF1A

**DOI:** 10.1101/2025.02.26.640415

**Authors:** Lu Rao, Wenxing Li, Yufeng Shen, Wendy K. Chung, Arne Gennerich

## Abstract

KIF1A is a neuron-specific kinesin motor responsible for intracellular transport along axons. Pathogenic *KIF1A* mutations cause KIF1A-associated neurological disorders (KAND), a spectrum of severe neurodevelopmental and neurodegenerative conditions. While individual *KIF1A* mutations have been studied, how different substitutions at the same residue affect motor function and disease progression remains unclear. Here, we systematically examine the molecular and clinical consequences of mutations at three key motor domain residues—R216, R254, and R307—using single-molecule motility assays and genotype-phenotype associations. Our findings reveal that mutations at R216 and R254 produce residue-specific effects, with some substitutions partially retaining function, whereas R307 mutations universally abolish motility, likely due to its critical role in microtubule binding. Mutant motor properties correlate with developmental outcomes, with R307 mutations linked to the most sever impairments. These results demonstrate that even single-residue substitutions can lead to distinct molecular and clinical phenotypes, highlighting the finely tuned mechanochemical properties of KIF1A. By establishing residue-specific genotype-phenotype relationships, this work provides fundamental insights into KAND pathogenesis and informs targeted therapeutic strategies.

## Introduction

KIF1A is a neuron-specific member of the kinesin-3 family, a group of microtubule (MT) plus-end-directed motor proteins essential for intracellular transport^1,2^. It plays a key role in nuclear migration in differentiating brain stem cells^3,4^ and facilitates the transport of synaptic precursors and dense core vesicles to axon terminals^5–10^. Over the past decade, more than 170 pathogenic *KIF1A* variants have been identified as the cause KIF1A-associated neurological disorders (KAND)^11^. KAND encompasses a broad spectrum of neurodevelopmental and neurodegenerative conditions, including progressive spastic paraplegias, microcephaly, encephalopathies, intellectual disability, autism, autonomic and peripheral neuropathy, optic nerve atrophy, and cerebral and cerebellar atrophy^11–53^.

KIF1A, like all kinesins, consists of a motor domain and a tail domain. The tail domain is highly divergent across kinesin family members, enabling functions such as oligomerization, cargo binding, and regulation^54^, while the motor domain is highly conserved and serves as the catalytic core^55^. This domain binds and hydrolyzes ATP to generate force and motion for microtubule-based transport. Its ATPase activity dependents on three key elements—the P-loop, switch-1 loop, and switch-2 loop^56,57^—which coordinate ATP-Mg binding and hydrolysis in a process stimulated by microtubule interaction^58,59^. In its auto-inhibited state, full-length KIF1A is monomeric^60^, with its tail domain folding back onto the motor domain to prevent dimerization^61^. Upon activation via phosphorylation^62^ or cargo binding^63^, KIF1A dimerizes and becomes highly processive^64^.

The severity and specific manifestations of KAND depend on the location of the mutation within the KIF1A heavy chain and the nature of the amino acid substitution, with most pathogenic variants residing in the KIF1A motor domain^11^ (**Fig. 1**). While several *KIF1A* missense mutations have been characterized at molecular^11,65–68^ or clinical level^11,69–72^, the effects of different amino acid substitutions at the same residue remain poorly understood. Establishing such connections is critical for elucidating disease mechanisms and guiding potential therapeutic strategies.

**Figure 1.**
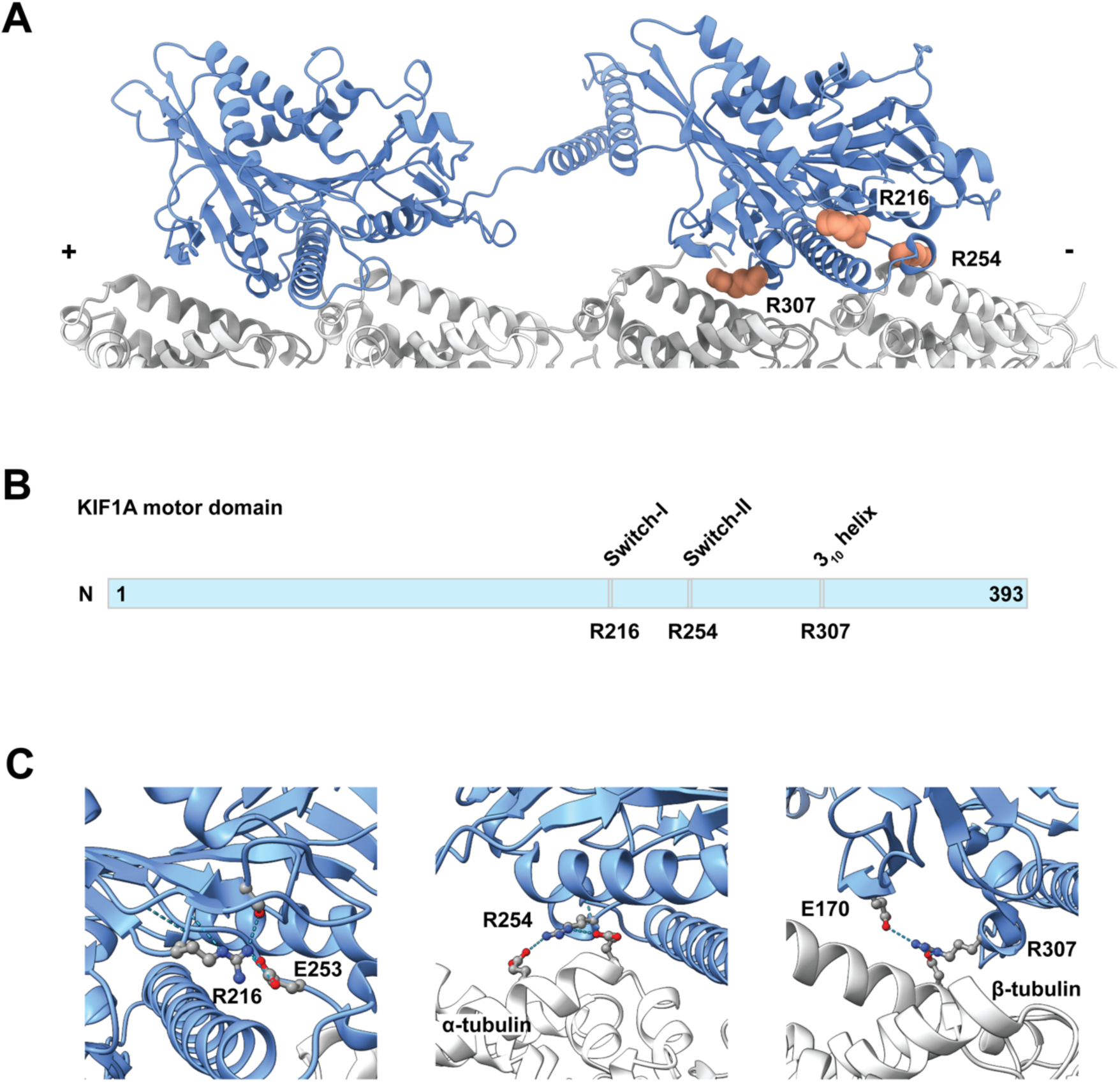
**(A)** Structure of a dimeric KIF1A bound on microtubules in the AMP-PnP state (PDB 8UTN)^65^. Residues R216, R254, and R307 in the trailing head are shown in pink in a space-filling representation. **(B)** Scheme of the KIF1A motor domain (amino acids 1-393) highlighting the locations of the three residues. **(C)** Close-up views of residue interactions: (Left) R216 interacts with S214 in the switch-I loop and E253 in the switch-2 loop. (Middle) R254 interacts with E414 and E420 in α-tubulin. (Right) R307 interacts intramolecularly with E170 and with D417 in β-tubulin.

Here, we examined the molecular and clinical consequences of three distinct KAND-associated amino acid substitutions at three different residues in the KIF1A motor domain: R216, R254, and R307. To investigate their effects, we used a tail-truncated, dimerizing KIF1A construct (393 amino acids) expressed in *E. coli*, which recapitulates the behavior of constitutively active full-length KIF1A in single-molecule assays^67^.

Our findings reveal that the magnitude of the biophysical alterations induced by these mutations correlates with the severity of clinical phenotypes. Specifically, differences in residue location and amino acid properties influence KIF1A’s motility defects and, consequently, the extent of neurodevelopmental impairment. These results establish a direct link between the biophysical consequences of residue-specific KIF1A mutations and their clinical impact, offering insights into disease mechanisms and potential therapeutic avenues.

## Results

### KIF1A mutants exhibit distinct single-molecule motility behaviors

To investigate how different amino acid substitutions at the same residue affect KIF1A function, we analyzed three conserved motor domain sites: R216, R254, and R307 (Fig. 1A). These residues are highly conserved or semi-conserved across kinesin families (Supp. Fig. 1). R216 is located at the center of the switch-1 loop, which regulates nucleotide exchange^57^. R254 is positioned at the end of the switch-2 loop, which transmits conformational changes critical for force generation^73^. R307 lies within the conserved YPRxS motif (where x represents D, E, or N), forming a unique 3_10_-helix adjacent to loop 12^66^ (Fig. 1B).

Previous studies have shown that disease-associated mutations in highly conserved motor domain residues often impair KIF1A function^11,66,67^. Here, we demonstrate that mutations at these three sites either abolish motility or significantly reduce velocity and processivity (Fig. 2). Moreover, different amino acid substitutions at the same site can lead to distinct molecular behaviors, emphasizing the role of residue-specific effects in determining motor dysfunction and clinical severity.

**Figure 2.**
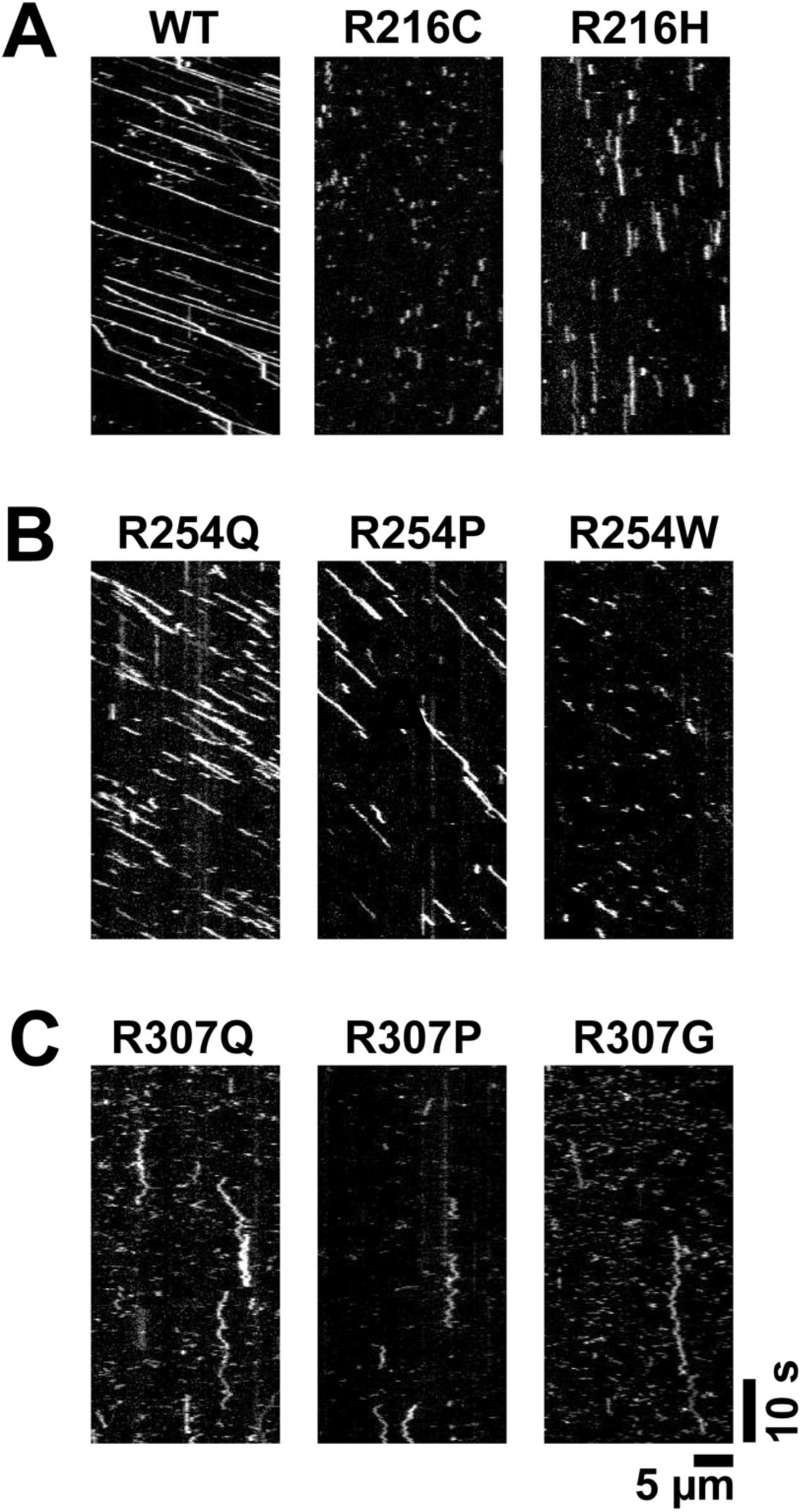
Representative kymographs of KIF1A mutants. **(A)** Wild-type (WT) KIF1A (left), R216C (middle) and R216H (right). **(B)** R254Q, R254P, and R254W. **(C)** R307Q, R307P, and R307G.

To probe these effects, we employed total internal reflection fluorescence (TIRF) microscopy to measure single-molecule motility using a tail-truncated, dimerizing KIF1A construct (residues 1-393). Engineered with a dimerizing leucine zipper to mimic the activated state of full-length KIF1A^67^, this construct exhibits a velocity of ∼2.3 µm/s and a run length of ∼15 µm (Fig. 2A, Table 1). We systematically introduced KAND-associated mutations at R216, R254, and R307 (Fig. 1A) and quantified their effects on velocity, processivity, and microtubule binding.

**Table 1.** Motility of KIF1A WT and mutants. The values represent median value with 95% confidence interval. WT: n=388; R216C: n=214; R216H: n=263; R254Q: n=304; R254P: n=354; R254W: n=235.

| | Velocity ($\mu\text{m/s}$) | Processivity ($\mu\text{m}$) | Dwell time (s) |
| --- | --- | --- | --- |
| WT | 2.34 [2.23, 2.36] | 14.5 [13.3, 15.6] | 6.4 [5.9, 6.8] |
| R216C | - | - | 0.97 [0.89, 1.04] |
| R216H | 0.096 [0.091, 0.107] | 0.46 [0.44, 0.49] | 4.7 [4.5, 5.2] |
| R254Q | 1.43 [1.40, 1.47] | 2.3 [2.0, 2.5] | 1.6 [1.5, 1.8] |
| R254P | 0.80 [0.78, 0.81] | 1.7 [1.4, 1.8] | 2.1 [1.9, 2.3] |
| R254W | 1.09 [1.05, 1.14] | 1.2 [1.1, 1.4] | 1.2 [1.0, 1.2] |
| R307Q | - | - | < 0.2 |
| R307P | - | - | < 0.2 |
| R307G | - | - | < 0.2 |

### R216 mutants impair KIF1A motility

R216, located within the switch-1 loop, plays a key role in KIF1A’s mechanochemical cycle by forming a salt bridge with the highly conserved E253 in the switch-2 loop (Fig. 1C)^74^. The switch-1 loop undergoes substantial conformational changes during the kinesin ATPase cycle, adopting a closed conformation in the ATP-bound state and an open conformation in the ADP-bound state^65^. It has been proposed that the R216–E253 interaction functions as a “backdoor” to facilitate efficient γ-phosphate release following ATP hydrolysis^74^. However, cryo-EM structures of KIF1A bound to microtubules in AMP-PnP (a non-hydrolysable ATP analog) and ADP states suggest that this interaction remains intact in both states^65^ (Fig. 1C, left; Suppl. Fig. 2A). This implies that the R216–E253 bond may break (“open the backdoor”) only when the motor domain detaches from microtubules, triggering γ-phosphate release.

Supporting this model, a recent structural study of KIF5B revealed that E236 (the equivalent of E253 in KIF1A) interacts with R203 (R216 in KIF1A) when bound to microtubules^75^. However, in the ADP state, when the motor domain is in solution, E236 instead interacts with T87 (T99 in KIF1A) of the P-loop^76^, a finding corroborated by another structural study^56^. The R216–E253 interaction likely prevents premature ADP release, as mutating E236 to alanine in KIF5B leads to a 100-fold increase in ADP release rate compared to wild-type^77^. Furthermore, mutations at E236 in KIF5B or R216 in KIF1A result in significantly slower ATP hydrolysis rates^78^, emphasizing the importance of this interaction in KIF1A’s ATPase cycle.

Mutations at R216 significantly impair KIF1A motility, though with distinct effects depending on the amino acid substitution (Fig. 2A, middle and right). The R216C mutation completely abolishes KIF1A motility along microtubules (Fig. 2A, middle), consistent with previous findings^24^. In contrast, R216H retains a low level of motility (Fig. 2A, left, Table 1), likely due to the partial preservation of local interactions by the charged histidine (His). Graph-based deep learning predictions^79^ suggest that while R216H does not interact with E253, it maintains interactions with the α4 helix, a region directly involved in microtubule binding of KIF1A^65^.

### R254 mutants retain motility along microtubules

R254 is a semi-conserved residue (Supp. Fig. 1) located at the end of the switch-2 loop. Its position suggests that conformational changes in the ATP-binding pocket may directly influence microtubule binding during the ATP hydrolysis cycle. Molecular modeling predicts that R254 in KIF1A (K237 in KIF5B) preferentially interacts with α-tubulin in ATP state compared to the ADP state^80,81^. The hypothesis is supported by cryo-EM structures of KIF1A bound to microtubules in either the AMP-PnP or ADP state^65^. In the AMP-PnP state, R254 interacts with two glutamate residues (E414 and E420) on α-tubulin (Fig. 1C, middle). However, in ADP state, R254 loses this interaction and instead forms an intramolecular interaction with D339 in H6 of KIF1A (Supp. Fig. 2B).

Despite its strong interactions with α-tubulin in ATP state, mutations at R254 do not abolish KIF1A’s motility (Fig. 2B), suggesting that R254 facilitates but is not essential for motor function. Consistent with this, a previous study reported that full-length KIF1A carrying the R254Q mutations retains motility^82^. This may explain why R254 is less conserved compared to core kinesin residues, such as those within the switch loops, which have more fundamental roles in motor activity.

Interestingly, different substitutions at R254 result in distinct single-molecule phenotypes. R254Q exhibits higher velocity and longer run length than R254P and R254W (Fig. 2B, Tab. 1), likely due to the chemical similarity between glutamine (Q) and arginine (R). While glutamine lacks arginine’s positive charge, it maintains a comparable size and hydrophilicity, making it more functionally compatible than proline or tryptophan. Structural predictions^79^ suggest that R254Q partially preserves interactions with E414 on α-tubulin in the AMP-PnP state (Supp. Fig. 3). In contrast, R254W disrupts the local environment due to the hydrophobicity of its aromatic ring, leading to steric clashes (Supp. Fig. 3). Meanwhile, R254P, with its much smaller side chain, does not participate in significant intramolecular or intermolecular interactions, and its effect likely stems from losing microtubule interactions.

### R307 mutants demonstrate only brief interactions or diffusion on microtubules

R307 is a highly conserved residue among kinesin families, located in the 3_10_-helix following loop 12^66^ (Fig. 1). Disease-associated mutations in this region, including P305L, Y306C, and R307P/Q/G, have been identified in patients with KAND^11^. Cryo-EM structures of dimeric KIF1A bound to microtubules in the presence of AMP-PnP indicate that R307 interacts with the H12 helix of β-tubulin^65^. Structural studies suggest that R278 in KIF5B (the equivalent to R307 in KIF1A) is stabilized by a salt bridge between its neighboring D279 and an arginine on the H8-S7 loop of β-tubulin, facilitating its interaction with the H12 helix^83^. This interaction is essential for kinesin’s microtubule-binding affinity.

Molecular modeling^80,81^ and a structural study^65^ (Fig. 1C, right; Suppl. Fig. 2C) further support R307 as a key contributor to microtubule binding in both the ATP and ADP states. A mutagenesis study on KIF5B revealed that replacing R278 (R307 in KIF1A) with alanine increases the motor’s Michaelis-Menton constant (K_m_) for microtubules more than 10-fold compared to wild-type kinesin, indicating a significant reduction in microtubule-binding affinity, while its microtubule-gliding velocity remained only slightly affected^84^. These findings suggest that mutations at this site do not abolish ATPase activity but instead severely impair microtubule attachment.

Mutations at R307 render KIF1A immobile, regardless of the substituting amino acid (Fig. 2C). The mutants interact only briefly with microtubules before dissociating, with occasional motor entering a diffusional state along the microtubule. A previous study also showed that R307P and R307Q mutants are immobile^85^. However, in that study, the mutants exhibited extensive diffusion along microtubules rather than brief interactions. The discrepancy likely arises from differences in buffer conditions: the previous study used BRB12, which has significantly lower ionic strength than BRB80, the buffer used here. Low ionic strength may enhance electrostatic interactions, promoting stronger motor-microtubule association.

The single-molecule behaviors of R307Q, R307P, and R307G are similar, with interaction durations shorter than 200 milliseconds. Unlike R254Q, where arginine-to-glutamine substitution partially preserves function, the R307 mutation does not retain motility. This underscores the importance of the salt bridge between R307 and β-tubulin in stabilizing microtubule binding.

### Mutant KIF1A motor properties correlate with clinical outcomes

To examine the relationship between KIF1A motor function and clinical severity, we analyzed Vineland Adaptive Behavior Scale (VABS) Adaptive Behavior Composite (ABC) scores— standardized measures used to assess adaptive behavior, daily living skills, and intellectual and developmental disabilities—in individuals carrying mutations in R216, R254, or R307. The cohort included 9, 28, and 9 individuals heterozygous for mutations at these residues, respectively. Mutant motor properties (velocity, processivity, and diffusion) were significantly associated with VABS ABC scores (**Table 2**). Individuals with R307 mutations exhibited markedly lower VABS ABC scores than those with R254 mutations (mean: 39 vs. 65, t-test, *p*=0.001) or R216 mutations (mean: 39 vs. 64, p=0.003). This aligns with our molecular data, which show that R307 mutants are largely immobile, whereas R216 and R254 mutants retain some mobility. Among R254 mutations, carriers of R254Q had marginally higher VABS ABC scores than individuals with other R254 mutations (mean: 76 vs. 58, p=0.04; **Table 3**). This is consistent with our single-molecule motility assays, which indicate that the R254Q mutant exhibits slightly higher velocities and run lengths compared to R216, R307, and other R254 variants.

**Table 2.** Univariate linear regression of key molecular phenotypes and VABS ABC.

| Variable | N | Mean (SD) | Beta (SE) | P |
| --- | --- | --- | --- | --- |
| Diffusion |  |  |  |  |
| No | 28 | 65.14 ± 11.07 | Reference | |
| Yes | 18 | 51.22 ± 19.12 | -13.92 (4.44) | 0.003 |
| Velocity | 46 |  | 14.18 (3.80) | 0.001 |
| Processivity | 46 |  | 11.80 (2.82) | <0.001 |
| Dwell time | 46 |  | 3.40 (1.74) | 0.058 |

**Table 3.**
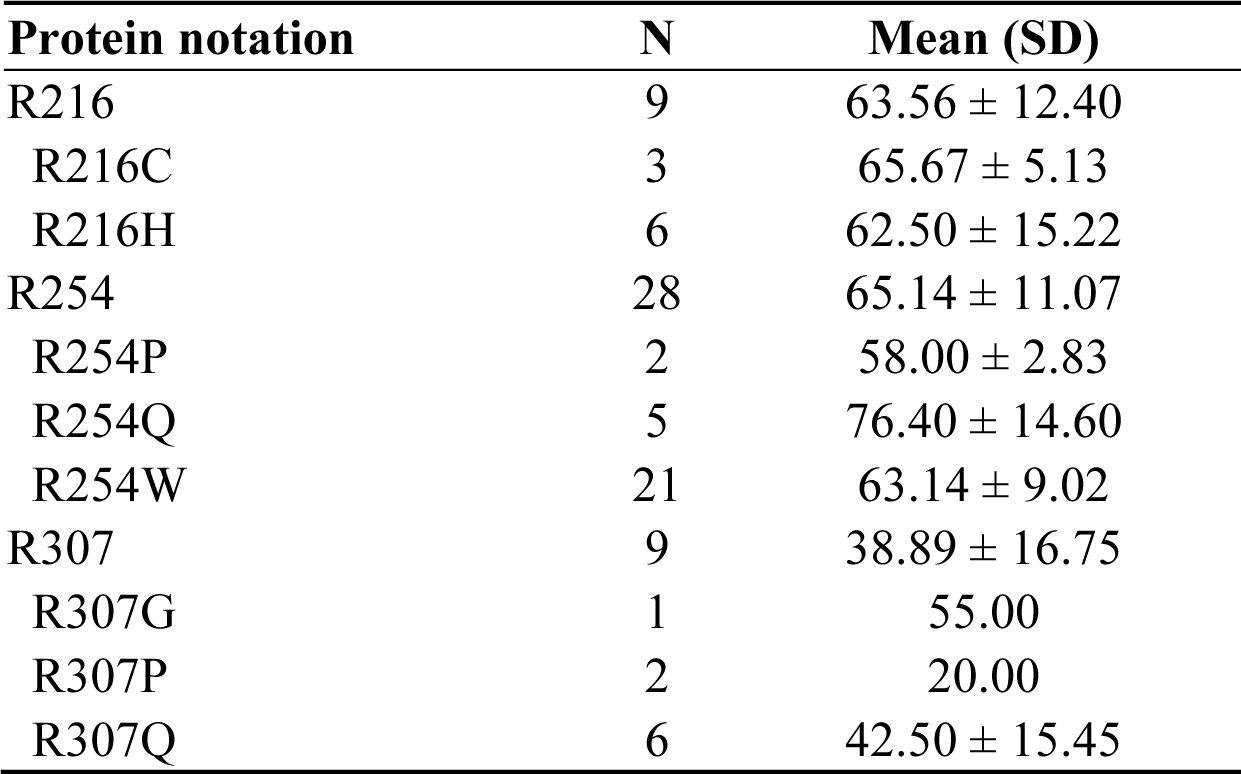
Summary of VABS ABC in patients with heterozygous *KIF1A* mutations.

| Protein notation | N | Mean (SD) |
| --- | --- | --- |
| R216 | 9 | 63.56 ± 12.40 |
| R216C | 3 | 65.67 ± 5.13 |
| R216H | 6 | 62.50 ± 15.22 |
| R254 | 28 | 65.14 ± 11.07 |
| R254P | 2 | 58.00 ± 2.83 |
| R254Q | 5 | 76.40 ± 14.60 |
| R254W | 21 | 63.14 ± 9.02 |
| R307 | 9 | 38.89 ± 16.75 |
| R307G | 1 | 55.00 |
| R307P | 2 | 20.00 |
| R307Q | 6 | 42.50 ± 15.45 |

These findings establish a direct link between KIF1A motility defects and clinical severity, demonstrating that even subtle differences in motor function can significantly impact neurodevelopmental outcomes.

## Discussion

KIF1A plays an essential role in intracellular transport by delivering synaptic vesicle precursors (SVPs) and dense-core vesicles along axons in neurons^5–10^. Given its importance in neuronal function, pathogenic mutations in *KIF1A* give rise to a broad spectrum of neurodevelopmental and neurodegenerative disorders known as KAND^11^. Understanding how specific mutations affect KIF1A’s function is key for deciphering the molecular mechanisms underlying these disorders and for informing therapeutic strategies. While previous studies have characterized the effects of individual KIF1A mutations at the single-molecule level, the impact of different amino acid substitutions at the same residue on motor function and disease progression has remained unexplored.

Here, we systematically examined three sets of KAND-associated mutations^11^ at distinct residues within the KIF1A motor domain—R216, R254, and R307—to determine how different substitutions at the same site impact KIF1A’s motility. Our findings reveal that the effects of these mutations vary significantly depending on their structural context and role in the motor’s mechanochemical cycle. Importantly, we show that the mobility properties of mutant KIF1A correlate with developmental outcomes in individuals carrying these mutations, providing molecular insights into clinical heterogeneity observed in KAND.

Mutations at R216, a key residue in the switch-1 loop, resulted in severe motility defects but exhibited mutation-specific differences. R216C completely abolished motility, whereas R216H retained a low level of movement, likely due to residual local interactions. Given that R216 forms a salt bridge with E253 in the switch-2 loop^74^, our findings support a model in which this interaction is essential for regulating ATP hydrolysis and microtubule attachment. The variation in motility among R216 substitutions suggest that partial functional compensation may occur when key structural interactions are preserved.

In contrast, R254 mutations, located at the end of the switch-2 loop, affected KIF1A function in a substitution-dependent manner but did not eliminate motility. R254Q, which retains some biochemical similarity to the wild-type arginine, exhibited higher velocity and longer run lengths than R254P or R254W. Structural predictions suggest that R254Q partially preserves interactions with α-tubulin, whereas R254W and R254P disrupt microtubule contacts through steric clashes or loss of stabilizing interactions. This suggest that R254 facilitates—but does not strictly dictate—KIF1A function, allowing for partial rescue depending on the nature of the substitution.

Unlike R216 and R254, mutations at R307, a highly conserved residue in the 310-helix following loop 12^66^, resulted in uniformly severe motility defects, regardless of substitution. All R307 mutants showed only brief interactions or diffusion along microtubules, with no sustained motility. Structural and modeling analyses indicate that R307 forms a critical interaction with the H12 helix of β-tubulin^65,80,81^, stabilizing microtubule attachment in both the ATP- and ADP-bound states. Disruption of this interaction significantly reduces KIF1A’s microtubule-binding affinity, which is consistent with previous findings in KIF5B^84^.

Consistent with our molecular findings, mutations at R307 are associated with the most severe developmental outcomes in KAND patients, likely dye ti the complete loss of motor function. Patients with R307 mutations exhibit profound motor impairments and cognitive deficits, reflecting the critical role of microtubule attachment in KIF1A-driven intracellular transport. These findings suggest that therapeutic strategies for R307-associated mutations may need to focus on compensatory mechanisms, such as enhancing microtubule binding through small-molecule stabilizers or gene therapy approaches aimed at restoring wild-type motor activity.

In contrast, individuals carrying the R254Q mutation exhibit relatively better developmental outcomes, aligning with its partial preservation of KIF1A motility. This suggests that small differences in biophysical properties, such as enhanced velocity and processivity, may translate into measurable clinical benefits. These findings raise the possibility that therapies aimed at fine-tuning motor activity—rather than fully restoring wild-type function—could be beneficial for specific *KIF1A* mutations. Given the mutation-dependent variability in disease severity, personalized therapeutic approaches tailored to individual genetic variants could be a promising strategy for KAND.

Together, our findings highlight the context-dependent effects of *KIF1A* mutations, where the same residue can have distinct impacts on motor function depending on the nature of the substitution and its role in the mechanochemical cycle. The mutation-specific differences at R216 and R254 contrast with the uniform loss of function seen at R307, emphasizing that KIF1A has undergone high evolutionary optimization for its distinct intracellular transport roles. These molecular insights provide a framework for understanding the clinical heterogeneity of KAND and underscore the need for mutation-specific therapeutic strategies. By establishing a direct link between residue-specific biophysical defects and their effects on motility, this study lays the groundwork for potential precision medicine approaches targeting KIF1A dysfunction. Future research should focus on leveraging this genotype-phenotype relationship to develop targeted interventions aimed at restoring KIF1A function in patients with KAND.

## Author Contributions

L.R. generated, expressed, and purified the constructs for the single-molecule studies. L.R. also designed and performed single-molecule experiments, collected and analyzed the data, and interpreted the results in collaboration with A.G. W.L. and Y.S. conducted and interpreted the statistical analysis of clinical data, which were collected by W.C. A.G. and W.C. conceived and coordinated the project. L.R, W.L.,Y.S., A.G. and W.C. wrote the manuscript. A.G. and W.C. secured funding.

## Acknowledgements

This work was supported by the National Institutes of Health Grants R01NS114636 (L. R., A.G. and W. C.) and R35GM149527 (Y.S.).

## Materials and Methods

### Plasmids and constructs

KIF1A mutants were generated from a previously described construct^11,65,67^ using Q5 mutagenesis (New England Biolabs, # E0554S). All the constructs were verified by Sanger sequencing (Genome Core Facility, Albert Einstein College of Medicine, Bronx, NY).

### *E. coli*-based KIF1A expression

All constructs were expressed in *E. coli*. Each plasmid was transformed into BL21-CodonPlus(DE3)-RIPL competent cells (Agilent Technologies, #230280). A single colony was picked and inoculated in 1 mL of terrific broth (TB) [10.1101/pdb.rec8620] containing 50 µg/mL carbenicillin and 50 µg/mL chloramphenicol. The culture was shaken at 37°C for 5 hours and then cooled to 16°C for 1 hour. Protein expression was induced by 0.1 mM IPTG overnight at 16°C. Cells were harvested by centrifugation at 3,000×g for 10 minutes at 4°C. The supernatant was discarded, and the pellet was fully resuspended in 5 mL of B-PER™ Complete Bacterial Protein Extraction Reagent (ThermoFisher Scientific, #89821) supplemented with 2 mM MgCl_2_, 1 mM EGTA, 1 mM DTT, 0.1 mM ATP, and 2 mM PMSF. The resuspension was flash-frozen in liquid nitrogen and stored at –80 °C.

### Purification of *E. coli*-expressed constructs

To purify *E. coli*-expressed protein, the frozen cell pellet was thawed at 37 °C and nutated at room temperature (RT) for 20 minutes to lyse the cells. For heterodimers, the two constructs were combined before nutation. The lysate was clarified by centrifugation at 80,000 rpm (260,000×g, *k*-factor=28) for 10 minutes in an TLA-110 rotor using a Beckman Tabletop Optima TLX Ultracentrifuge. The supernatant was passed through 500 μL of Ni-NTA Roche cOmplete™ His-Tag purification resin (Millipore Sigma, #5893682001). The resin was washed with 4 mL of wash buffer (50 mM HEPES, pH 7.2, 300 mM KCl, 2 mM MgCl_2_, 1 mM EGTA, 1 mM DTT, 1 mM PMSF, 0.1 mM ATP, 0.1% (w/v) Pluronic F-127, 10% glycerol). To label the SNAPf-tag, SNAP-Cell® TMR-Star ligand (New England Biolabs, # S9105S) was added to the resin to a final concentration of 10 µM before elution, and the resin was incubated at RT for 20 minutes. The resin was then washed with 6 mL of wash buffer, and the protein was eluted using elution buffer (50 mM HEPES, pH7.2, 150 mM KCl, 2 mM MgCl_2_, 1 mM EGTA, 1 mM DTT, 1 mM PMSF, 0.1 mM ATP, 0.1% (w/v) Pluronic F-127, 10% glycerol, 150 mM imidazole). The elute was flash frozen and stored at –80 °C.

### Microtubule polymerization

To polymerize microtubules, 2 µL of 10 mg/mL tubulin (Cytoskeleton, #T240-B) was mixed with 2 µL of 1 mg/mL biotinylated tubulin (Cytoskeleton, #T333P-A), 1 µL of 1 mg/mL HiLyte488-labeled tubulin (Cytoskeleton, #TL488M-A), and 1 µL of 10 mM GTP. The mixture was incubated at 37°C for 20 minutes, followed by the addition of 0.6 µL of 0.2 mM paclitaxel in DMSO. Incubation continued for another 20 minutes. The polymerized microtubules were then carefully layered on top of 100 µL of a glycerol cushion (80 mM PIPES, pH 6.8, 2 mM MgCl_2_, 1 mM EGTA, 60% (v/v) glycerol, 1 mM DTT, 10 µM paclitaxel) in a 230-µL TLA100 tube (Beckman Coulter, #343775) and centrifuged at 80,000×rpm (250,000×g, *k*-factor=10) for 5 minutes at RT using a Beckman Tabletop Optima TLX Ultracentrifuge. The supernatant was carefully removed, and the pellet was resuspended in 11 µL of BRB80G10 (80 mM PIPES, pH 6.8, 2 mM MgCl_2_, 1 mM EGTA, 10% (v/v) glycerol, 1 mM DTT, 10 µM paclitaxel). The microtubule solution was stored at RT in the dark for further use. For the microtubule-binding and -release assay, only 5 µl of 10 mg/mL unlabeled tubulin was used for polymerization.

### Microtubule-binding and -release (MTBR) assay

An MT-binding and -release (MTBR) assay was performed to remove inactive motors prior to single-molecule TIRF assays. Fifty microliter of each protein construct was buffer-exchanged into a low salt buffer (30 mM HEPES, pH 7.2, 50 mM KCl, 2 mM MgCl_2_, 1 mM EGTA, 1 mM DTT, 1 mM AMP-PNP, 10 µM paclitaxel, 0.1% (w/v) Pluronic F-127, and 10% glycerol) using a 0.5-mL Zeba™ spin desalting column (7-kDa MWCO) (ThermoFisher Scientific, #89882). After buffer exchange, the solution was warmed to RT and 3 μL of 5 mg/mL paclitaxel-stabilized microtubules was added. The mixture was gently mixed and layered over a 100 μL glycerol cushion (80 mM PIPES, pH 6.8, 2 mM MgCl2, 1 mM EGTA, 1 mM DTT, 10 µM paclitaxel, and 60% glycerol). The sample was centrifuged at 45,000 rpm (80,000×g, *k*-factor=33) for 10 minutes at RT using a TLA-100 rotor in a Beckman Tabletop Optima TLX Ultracentrifuge. The supernatant was carefully removed, and the pellet was resuspended in 50 μL high-salt release buffer (30 mM HEPES, pH 7.2, 300 mM KCl, 2 mM MgCl_2_, 1 mM EGTA, 1 mM DTT, 10 μM paclitaxel, 1 mM ATP, 0.1% (w/v) Pluronic F-127, and 10% glycerol). Microtubules were then removed by centrifugation at 40,000 rpm (60,000×g, *k*-factor=41) for 5 minutes at RT. The final supernatant, containing the active motors, was aliquoted, flash-frozen in liquid nitrogen, and stored at –80 °C.

### Total internal reflection fluorescence (TIRF) assay

#### Flow chamber preparation

A flow chamber was assembled using a glass slide (Fisher Scientific, #12-550-123) and an ethanol-cleaned coverslip (Zeiss, #474030-9000-000) with two thin strips of parafilm as spacers. To functionalize the surface, 10 µL of 0.5 mg/ml BSA-biotin (ThermoScientific, #29130) was introduced into the chamber and incubated for 10 minutes. The chamber was then washed with 2×20 µL blocking buffer (80 mM PIPES, pH 6.8, 2 mM MgCl2, 1 mM EGTA, 10 µM paclitaxel, 1% (w/v) Pluronic F-127, 2 mg/mL BSA, 1 mg/mL α-casein) and incubated for 30 minutes to block the surface. Next, 10 µL of 0.25 mg/ml streptavidin (Promega, #Z7041) was introduced into the chamber and incubated for 10 minutes, followed by another wash with 2×20 µL blocking buffer. Ten microliters of 0.02 mg/mL biotin-labeled microtubules in the blocking buffer was then introduced and incubated for 1 minute. The chamber was washed again with 2×20 µL blocking buffer and stored in a humid chamber until use.

#### Sample preparation

The motor solution was diluted to the appropriate concentration, and 1 µL of the diluted motor solution was added to 50 µL of motility buffer (80 mM PIPES, pH 7.2, 2 mM MgCl_2_, 1 mM EGTA, 10 µM paclitaxel, 0.5% (w/v) Pluronic F-127, 5 mg/mL BSA, 1 mg/mL α-casein, 2 mM ATP, 2 mM biotin, gloxy oxygen scavenger system). Two times 20 µL of the solution was introduced into the chamber, which was then sealed with vacuum grease.

#### Data acquisition

Images were acquired using BioVis software (BioVision Technologies) with an acquisition time of 200 ms per frame and 600 frames per movie.

#### Data analysis

Kymographs were generated using Fiji^87^, and the velocity and run length were analyzed using a home-built MATLAB program.

### Graphic visualization

The molecular structures in the figures were visualized using UCSF ChimeraX^88^ or PyMOL (version 3.1).

### Clinical studies

All studies were approved by the Institutional Review Boards at Columbia University and Boston Children’s Hospital. Informed consent was obtained from all patients or their guardians. Patient recruited, genetic diagnosis verification, and clinical characterization were conducted as previously described^11,71^. For individuals with multiple Vineland score assessments, the most recent score was used for statistical analysis.

