## Supplementary Figures for "Distinct Clinical Phenotypes in KIF1A-Associated Neurological Disorders Result from Different Amino Acid Substitutions at the Same Residue in KIF1A"

```
kif21b      ----------MAGQGDCCVKVAVRIRPQLSKEKIEGCHICTSVTPGE--------PQVLL   42
kif1a       -------------MAGASVKVAVRVRPFNSREMSRDSKCIIQMSGST---TTIVNPKQPK   44
kif5b       ----------MADLAECNIKVMCRFRPLNESEVNRGDKYIAK----------FQGEDTVV   40
kif3a       ---MP-INKSEKPESCDNVKVVVRCRPLNEREKSMCYKQAVSVDEMRGTITVHKTDS-SN   55
kif11       MASQPNSSAKKKEEKGKNIQVVVRCRPFNLAERKASAHSIVECDPVRKEVSVRTGGLADK   60
                              ::*  * **    *     :   .

kif21b      GKDKAFTYDFVFDLDT--------WQEQIYSTCVSKLIEGCFEGYNATVLAYGQTGAGKT   94
kif1a       ETPKSFSFDYSYWSHTSPEDINYASQKQVYRDIGEEMLQHAFEGYNVCIFAYGQTGAGKS   104
kif5b       IASKPYAFDRVFQSST--------SQEQVYNDCAKKIVKDVLEGYNGTIFAYGQTSSGKT   92
kif3a       EPPKTFTFDTVFGPES--------KQLDVYNLTARPIIDSVLEGYNGTIFAYGQTGTGKT   107
kif11       SSRKTYTFDMVFGAST--------KQIDVYRSVVCPILDEVIMGYNCTIFAYGQTGTGKT   112
               * :::*  :   :         * ::*      ::.  : ***  ::*****.:**:

kif21b      YTMGTGFDMA-----TSEEEQGIIPRAIAHLFGGIAERKRRAQEQGVAGPEFKVSAQFLE   149
kif1a       YTMMGKQEK---------DQQGIIPQLCEDLFSRINDTT-------NDNMSYSVEVSYME   148
kif5b       HTMEGKLHDP--------EGMGIIPRIVQDIFNYIYSM--------DENLEFHIKVSYFE   136
kif3a       FTMEGVRAIP--------ELRGIIPNSFAHIFGHIAKA--------EGDTRFLVRVSYLE   151
kif11       FTMEGERSPNEEYTWEEDPLAGIIPRTLHQIFEKLT----------DNGTEFSVKVSLLE   162
            .**                  ****.   .:*  :             .  : : .. :*

kif21b      LYNEEILDLFDSTRDPDTRHRRSNIKIHEDANGGIYTTGVTSRLIHSQEELIQCLKQGAL   209
kif1a       IYCERVRDLLNPKNKGN-----LRV--REHPLLGPYVEDLSKLAVTSYNDIQDLMDSGNK   201
kif5b       IYLDKIRDLLDVSKTN------LSV--HEDKNRVPYVKGCTERFVCSPDEVMDTIDEGKS   188
kif3a       IYNEEVRDLLGKDQT--QR---LEV--KERPDVGVYIKDLSAYVVNNADDMDRIMTLGHK   204
kif11       IYNEELFDLLNPSSDVSER---LQMFDDPRNKRGVIIKGLEEITVHNKDEVYQILEKGAA   219
            :* :.: **:.             :             .     : . :::   :  *

kif21b      SRTTASTQMNVQSSRSHAIFTIHLCQMRMCTQPDLVNEAVTGLPDGTPPSSEYETLTAKF   269
kif1a       ARTVAATNMNETSSRSHAVFNIIFTQKRHDAETNI-----------------TTEKVSKI   244
kif5b       NRHVAVTNMNEHSSRSHSIFLINVKQENTQT---------------------EQKLSGKL   227
kif3a       NRSVGATNMNEHSSRSHAIFTITIECSEKGIDGN------------------MHVRMGKL   246
kif11       KRTTAATLMNAYSSRSHSVFSVTIHMKETTIDGE------------------ELVKIGKL   261
             * .. * **  *****::* : .   .                             .*:

kif21b      HFVDLAGSERLKRTGATGERAKEGISINCGLLALGNVISALGDQ---------SKKVVHV   320
kif1a       SLVDLAGSERADSTGAKGTRLKEGANINKSLTTLGKVISALAEMDSGPNKNKKKKTDFI   304
kif5b       YLVDLAGSEKVSKTGAEGAVLDEAKNINKSLSALGNVISALAEG------------STYV   275
kif3a       HLVDLAGSERQAKTGATGQRLKEATKINLSLSTLGNVISALVDG-----------KSTHV   295
kif11       NLVDLAGSENIGRSGAVDKRAREAGNINQSLLTLGRVITALVE------------RTPHV   309
             :*******.   :** .    *. .** .* :**.**:** :               .:

kif21b      PYRDSKLTRLLQDSLGGNSQTIMIACVSPSDRDFMETLNTLKYANRARNIKNKVVVNQDK   380
kif1a       PYRDSVLTWLLRENLGGNSRTAMVAALSPADINYDETLSTLRYADRAKQIRCNAVINED-   363
kif5b       PYRDSKMTRILQDSLGGNCRTTIVICCSPSSYNESETKSTLLFGQRAKTIKNTVCVNVEL   335
kif3a       PYRNSKLTRLLQDSLGGNSKTMMCANIGPADYNYDETISTLRYANRAKNIKNKARINED-   354
kif11       PYRESKLTRILQDSLGGRTRTSIIATISPASLNLEETLSTLEYAHRAKNILNKPEVNQKL   369
            ***:* :* :*::.***. :* :    .*:. :  ** .** :..**: *  .  :* .
```

**Supplemental Figure 1.** Multiple sequence alignment of selected kinesins generated using Clustal Omega[1]. KIF1A residues R216, R254, and R307 are highlighted in red.

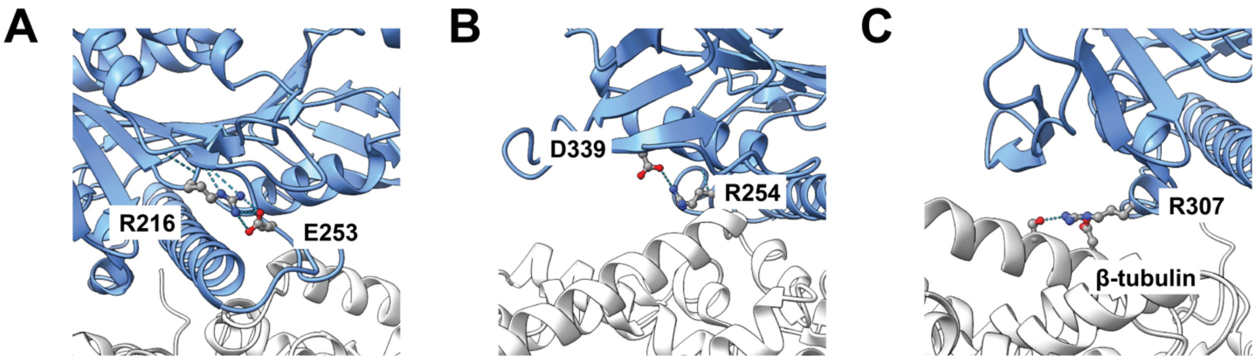


**Supplemental Figure 2.** Local interactions of KIF1A residues R216, R254, and R307 in the ADP state when bound to microtubules (PDB 8UTR)[2]. **(A)** R216 interacts with E253. **(B)** R254 interacts with D339. **(C)** R307 interacts with S413 and D417 in β-tubulin.

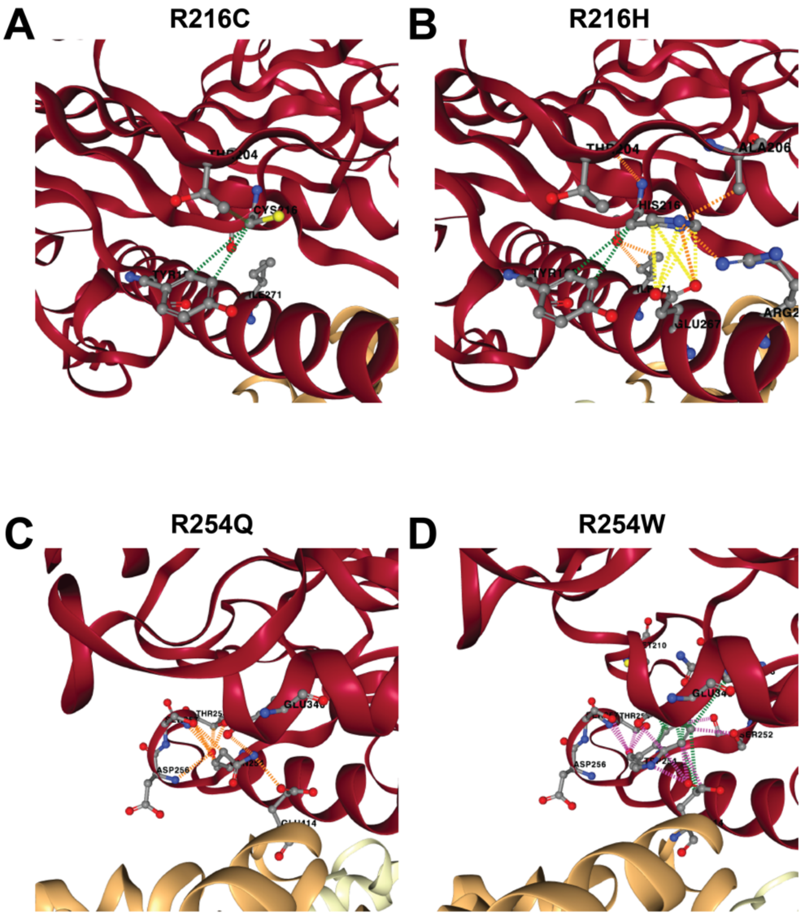


**Supplemental Figure 3.** Structural prediction of KIF1A mutants using graph-based deep learning.

1. Madeira, F., Madhusoodanan, N., Lee, J., Eusebi, A., Niewielska, A., Tivey, A.R.N., Lopez, R. & Butcher, S. The EMBL-EBI Job Dispatcher sequence analysis tools framework in 2024. *Nucleic Acids Res* **52**, W521-W525 (2024).
2. Benoit, M., Rao, L., Asenjo, A.B., Gennerich, A. & Sosa, H. Cryo-EM unveils kinesin KIF1A's processivity mechanism and the impact of its pathogenic variant P305L. *Nat Commun* **15**, 5530 (2024).
